# Dual roles for nucleus accumbens core dopamine D1-expressing neurons projecting to the substantia nigra pars reticulata in limbic and motor control

**DOI:** 10.1101/2023.03.25.534237

**Authors:** Suthinee Attachaipanich, Takaaki Ozawa, Tom Macpherson, Takatoshi Hikida

## Abstract

The nucleus accumbens (NAc) is a critical component of a limbic basal ganglia circuit that is thought to play an important role in decision-making and the processing of rewarding stimuli. As part of this circuit, dopamine D1 receptor-expressing medium spiny neurons (D1-MSNs) of the NAc core are known to send a major projection to the substantia nigra pars reticulata (SNr). However, the functional role of this SNr-projecting NAc D1-MSNs (NAc^D1-MSN^-SNr) pathway is still largely uncharacterized. Moreover, as the SNr is thought to belong to both limbic and motor information processing basal ganglia loops, it is possible that the NAc^D1-MSN^-SNr pathway may be able to influence both limbic and motor functions. In this study we investigated the effect of optogenetic activation of the NAc^D1-MSN^-SNr pathway on reward-learning and locomotor behavior. Stimulation of the axon terminals of NAc core D1-MSNs in the SNr induced a preference for a laser-paired location, self-stimulation via a laser-paired lever, and augmented instrumental responding for a liquid reward-paired lever. Additionally, stimulation was observed to increase locomotor behavior when delivered bilaterally and induced contralateral turning behavior when delivered unilaterally. These findings indicate that the NAc^D1-MSN^-SNr pathway is able to control both reward learning and motor behaviors.

## 1. Introduction

The nucleus accumbens (NAc) is a key component of the brain’s reward and motivation system and is a major input nucleus of the basal ganglia, a group of nuclei that are important for learning and motor control (Floresco, 2015; Macpherson and Hikida, 2019; Salgado and Kaplitt, 2015). Classically, the NAc has been delineated into at least two subregions, the central core (NAc core) and surrounding shell (NAc shell), based upon the expression of various histochemical markers, as well as topographically organized afferent inputs (Groenewegen et al., 1999; Jongen-Rêlo et al., 1994; Ma et al., 2020; Voorn et al., 2004; Záborszky et al., 1985; Zahm, 1999). Within both the NAc core and shell, medium spiny neurons (MSNs), the major neuron type, are typically subdivided into two subpopulations of roughly equal size according to their expression of dopamine receptors: dopamine D1 and D2 receptor-expressing MSNs (D1- and D2-MSNs) (Gerfen et al., 1990; Tepper and Bolam, 2004). While NAc D1-MSNs have been implicated in reward learning and the attribution of motivational salience (Hikida et al., 2010; Lobo et al., 2010; Macpherson et al., 2014; Macpherson and Hikida, 2018), D2-MSNs have been indicated to play an important role in aversive learning, stimulus discrimination, and behavioral flexibility (Hikida et al., 2010; Yawata et al., 2012; Macpherson et al., 2016; Iino et al., 2020; Macpherson et al., 2022). Additionally, it has been reported that NAc D1- and D2-MSNs may also contribute to the control of locomotor activity. Intra-NAc infusions of both D1 and D2 receptor agonists have been reported to increase locomotion activity in rats, while intra-NAc infusion of D1 and D2 antagonists reduced locomotor activity (Dreher and Jackson, 1989; Plaznik et al., 1989). In partial agreement with these studies, more recently, chemogenetic activation of NAc D1-MSNs was demonstrated to increase, while activation of NAc D2-MSNs to decrease, wheel running and locomotor activity in an open field arena (Zhu et al., 2016). These studies indicate that NAc D1- and D2-MSNs may contribute to the functional control of limbic as well as locomotor processing.

In the NAc core, D2-MSNs largely project to the ventral pallidum (VP), while NAc D1-MSNs are known to send equal projections to both the VP (NAc^D1-MSN^-VP) and the substantia nigra pars reticulata (SNr) (NAc^D1-MSN^-SNr) (Heimer et al., 1991; Kupchik et al., 2015; Kupchik and Kalivas, 2017; Matsui and Alvarez, 2018; Pardo-Garcia et al., 2019; Robertson and Jian, 1995). While the NAc^D1-MSN^-VP pathway has been established to play an important role in cocaine addiction related behaviors, including cue-induced reinstatement of cocaine-seeking and cocaine-induced locomotor sensitization (Stefanik et al., 2013; Creed et al., 2016; Pardo-Garcia et al., 2019), little is currently known about the functional role of the NAc^D1-MSN^-SNr pathway. Classically, NAc D1-MSN projections to the SNr have been suggested to form an integral part of a limbic information processing basal ganglia loop circuit beginning in the medial prefrontal cortex (mPFC) and connecting the NAc, the VP, the SNr, and the thalamus, before returning to the mPFC (Alexander et al., 1986; Foster et al., 2021; Haber, 2003; Macpherson et al., 2021; Parent and Hazrati, 1995). In this limbic circuit, activation of NAc D1-MSNs is hypothesized to drive reinforcement; however, this proposed role is yet to be empirically tested (Balleine, 2019; Macpherson et al., 2021; Peak et al., 2019). Interestingly, recent evidence has indicated that the NAc^D1-MSN^-SNr pathway may also be able to influence motor behavior. Indeed, optogenetic activation of NAc D1-MSNs axon terminals in the SNr were reported to increase activity not only in the mPFC, but also in the motor cortex (M1) (Aoki et al., 2019). While this study highlights the potential for the NAc^D1-MSN^-SNr pathway to exert an influence over motor areas such as the M1, it has yet to be established whether activation of this pathway results in changes in motor behavior.

Here we used activation optogenetic activation of the axon terminals of NAc core D1-MSNs in the SNr to investigate the contribution of the NAc^D1-MSN^-SNr pathway to limbic and motor functions. We reveal that stimulation of this NAc-to-SNr pathway was able to induce a strong reinforcing effect in place preference and self-stimulation tasks and could augment instrumental responding for a liquid reward. Further, we demonstrate that optogenetic activation of the NAc^D1-MSN^-SNr pathway resulted in increased motor activity in an open field arena. These findings indicate that the NAc^D1-MSN^-SNr pathway is able to play a dual role in controlling limbic and motor functions, and highlights the potential of targeting this circuit for interventions for clinical conditions associated with limbic or motor impairments.

## 2. Material and Methods

### 2.1 Animals

Group-housed male D1-Cre transgenic mice (FK150Gsat, Jackson Laboratory, USA) aged 8-12 weeks old and on a C57BL/6 background, as well as their wildtype counterparts, were used for experiments. Animals were housed in groups of 2-3 and were maintained on a 12-h light/dark cycle (lights on at 8:00 a.m.) with the temperature controlled to 24 ± 2 ºC in a humidity of 50 ± 5%. Behavioral experiments were conducted during the light period. Mice were provided access to water and standard lab chow *ad libitum*, except for during touchscreen operant chamber experiments during which time mice were food restricted to maintain motivation to instrumentally respond (see 2.3.2). All animal experiments conformed to the guidelines of the National Institutes of Health experimental procedures and were approved by the ethical committee of [Author’s University].

### 2.2 Stereotaxic Virus Injection and optical cannula Implantation

Following anesthesia (90 mg/kg Ketamine and 20 mg/kg Xylazine, i.p. injection), mice were positioned in a stereotaxic apparatus. A midline incision was made down the scalp and a craniotomy was made using a dental drill. Injections were carried out using graduated pipettes with a tip diameter of 10-15 μm. Mice were injected into the NAc core (Bregma coordinates: anterior/posterior, +1.2 mm; medial/lateral, ± 1.35 mm; dorsal/ventral, 3.75 mm, 250 nl/site at a rate of 100 nl/min) bilaterally with an adeno-associated virus (AAV) virus expressing channelrhodopsin-2 (ChR2) under a FLEX cre-switch promotor (AAV2-Ef1a-FLEX-hChR2(H134R)-EYFP), or an optically-inactive control virus (AAV2-Ef1a-FLEX-EYFP) at concentrations of 4.4 × 10^12^ virus molecules/ml and 5.3 × 10^12^ virus molecules/ml, respectively. Following infusion, the needle was kept at the injection site for 10 min to allow for diffusion of the virus and then slowly withdrawn. For robust viral expression, viral infusions occurred a minimum of 3-4 weeks before behavioral training. For behavioral experiments, chronically implantable optic fibers (200-μm core 0.22 N.A., Thorlabs, Newton, NJ, USA) threaded through ceramic zirconia ferrules were implanted bilaterally into the medial SNr (Bregma coordinates: anterior/posterior, -3.3 mm; medial/lateral, ± 1.0 mm; dorsal/ventral, 4.5 mm). Finally, three skull screws were implanted 1 mm into the skull surrounding the optic fibers and the whole skull was secured using dental cement.

For both ChR2-expressing D1-Cre mice and EYFP-expressing control mice, a 20 Hz stimulation protocol was used: 473 nm light, 8-10 mW, 10 ms pulse, delivered by DPSS lasers, which were controlled using a pulse generator.

### 2.3 Behavioral testing

#### 2.3.1 Real-time place preference test (rt-PP)

rt-PP was conducted in a white rectangular box divided lengthways down the center into two equal-sized rectangular chambers (W:15cm x L:20cm x H:25cm). Each chamber contained different contextual cues; one chamber had green triangles on the walls, while the other chamber had blue dots on the walls.

The rt-PP task was composed of two stages: (1) a pre-conditioning test (pre-test) and (2) test sessions (laser-test). During the pre-test, mice were able to freely explore the entire apparatus for 15 min and the time spent in each chamber was measured to check for any bias to either side by using automated video tracking software (EthoVision XT 16, Noldus). Next, the laser-test sessions were performed using an unbiased experimental design across 3 consecutive days for 20 min each day. When an animal entered into one chamber the laser was delivered (473 nm light, 8-10 mW, 10 ms pulse) for the duration that the animal stayed in the chamber, whereas when they entered the other chamber no stimulation was delivered. The chamber (triangle or dot walls) paired with laser stimulation was randomized between animals. The time spent in the laser-paired chamber was averaged across the three test sessions and then compared with the time spent in the non-laser-paired chamber to assess the mouse’s preference.

#### 2.3.2 Operant chamber tests of reinforcement

Operant tasks were conducted in trapezoidal Bussey-Saksida touchscreen operant chambers (Lafayette instrument, IN, USA) housed within a light- and sound-attenuating cubicles. In each chamber, the front touchscreen was divided into two touch response panels (70×75 mm^2^ spaced, 5 mm apart, 16 mm above the floor) and a liquid delivery magazine was placed at the back end of the chamber. Self-stimulation tests were controlled by ABET II and Whisker Server software (Lafayette instrument, IN, USA), and laser delivery in the chambers was controlled by Radiant v2 software (Plexon Inc, TX, USA).

##### 2.3.2.1 Two-choice optogenetic self-stimulation task

Mice were first food restricted until they reached 85-90% of their free-feeding weight (approx. 7 days) in order to increase their motivation to produce instrumental behavioral responses. Then, in four consecutive daily sessions, mice were trained under a fixed ratio 1 (FR1: 1 response produces the outcome) schedule to instrumentally respond at a touch panel paired with the delivery of laser stimulation (S+: 473 nm, 8-10 mW, 20Hz pulse for 30 sec) or a touch panel paired with delivery of no outcome (S-). The spatial (left/right) location of the S+ response panel was counterbalanced across mice. The outcome of each trial was followed by a 10-sec intertrial interval (ITI). Each session lasted 60 mins or until mice had completed 60 trials. The number of touch responses for the S+ and S-panels, as well as the latencies to make responses, were recorded in each session to assess the potential reinforcing effect of laser stimulation.

##### 2.3.2.2 Two-choice task with optogenetic stimulation paired with a liquid reinforcer

Next mice were trained to instrumentally respond at the same two response panels for delivery of a sucrose liquid reward (7µl of 10% sucrose diluted in water); however, while a response at one panel (previously paired with laser delivery) was additionally paired with delivery of laser stimulation (S+: 473 nm, 8-10 mW, 20Hz pulse for 30 sec), a response at the other panel (previously associated with no outcome) delivered just the liquid reward alone (S-). Each outcome was followed by a 10-sec ITI. The location of the S+ panel was counterbalanced across previously S+ and S-panels in the prior self-stimulation task. No main effect or interaction of previous panel location was observed so data were grouped together.

Animals were trained on consecutive days using a previously described schedule of reinforcement with minor modifications (Robinson et al., 2014; Soares-Cunha et al., 2022), as follows: FR1 schedule for 5 days, FR4 schedule for 1 day, random-ratio 4 (RR4) schedule (a random number of responses between 1-4 produces the outcome) for 4 days, and finally RR6 for 4 days each. As previous, each session lasted 60 mins or until 60 trials had been completed. The number of touch responses for the S+ (sucrose & laser) and the S-(sucrose alone), as well as the latencies to respond and collect the liquid rewards, was measured for each session.

#### 2.3.3 Open field test of motor activity (OFT)

For bilateral stimulation tests, mice were placed in a grey cylindrical (42 cm diameter, 42 cm height) open field apparatus. Patch cords were attached to bilateral fiber optic cannulae and suspended above the animal so that they could freely move to all areas of the apparatus. Animals were allowed to freely explore the entire arena and habituate to the apparatus for three min before the start of testing. The test consisted of a 12 min long session divided into four alternating 3 min trials during which bilateral laser stimulation was either OFF or ON (OFF-ON-OFF-ON), according to a previously described method (Tye et al., 2013). During ON trials, photostimulation was delivered according to the protocol described in 2.2. Total distance moved, velocity, and speed were automatically recorded by video tracking software (EthoVision XT 16, Noldus).

Unilateral stimulation was performed as described above, with the exception that only one patch cable was attached to either the contralateral or ipsilateral fiber optic cannulae. Controlateral and ipsilateral tests were performed on consecutive days with the order randomized across mice. Body rotations (>180° turn) were automatically recorded using video tracking software (EthoVision XT 16, Noldus).

### 2.3 Histological analysis

After behavioral experiments were completed, mice were anesthetized with 90 mg/kg Ketamine and 20 mg/kg Xylazine and transcardially perfused with 0.1 M phosphate buffered saline (PBS) for 2 minutes followed by 4% paraformaldehyde (PFA) (Nacalai Tesque Inc, Kyoto, Japan) in 1x PBS for 5 minutes at a 10 ml/min flow rate. Brains were removed and postfixed overnight in 4% PFA, then placed in 7.5%, 15%, and 30% sucrose in 1x PBS solutions at 4°C until the brains sank in the solution at each stage. Brains were embedded and frozen completely in Optimal Cutting Temperature (O.C.T.) compound (Sakura Finetek, Osaka, Japan). Then, the brain tissue was attached to a circular cryostat block and sectioned on a cryostat (Leica CM1860, Leica, Wetzlar, Germany) into 40-μm-thick slices at -17 to -20°C. Coronal brain slices (40 μm) were stored in PBS solution at 4°C. For immunohistochemical staining, each brain section was treated with a blocking solution (5% Bovine serum albumin (Nacalai Tesque Inc, Kyoto, Japan) in 1x PBS) for 1 h at room temperature and washed three times in 1x PBS. After rinsing in 1x PBS, the slices were incubated with anti-green fluorescent protein rabbit IgG primary antibody (1:1000; Molecular Probes, OR, USA) and anti-Tyrosine Hydroxylase (TH) antibody (1:500; EMD Millipore, USA) in 1x PBS with 0.3% Triton-X (Nacalai Tesque Inc, Kyoto, Japan) (PBST) overnight at 4°C. All brain sections were washed three times for 10 min in PBS, then stained with Alexa Fluor 488 or 555 goat anti-rabbit IgG secondary antibody (1:500; Thermo Fisher Scientific, MA, USA) in 1x PBST for 1 h at room temperature. After being washed again three times in 1x PBS for 10 min, the sections were mounted using Fluoroshield mounting medium containing DAPI (Abcam, Cambridge, UK) and observed using a KEYENCE BZ-X800E All-in-one Fluorescence Microscope (Keyence, Osaka, Japan).

### 2.4 Statistical Analyses

All experimental data was plotted as the mean ± SEM using Prism v8.0 software (GraphPad Software Inc, CA, USA). Results from all statistical analyses are shown in Supplementary table 1. The rt-PP and OFT (bilateral stimulation) data were analyzed using two-way repeated measures ANOVAs with virus (ChR2 vs EYFP) as a between-subjects factor, laser (OFF vs ON) as a within subjects factor, and time spent in the laser-paired chamber minus time spent in the non-laser paired chamber (for RT-PP), velocity (for OFT), or distance travelled (for OFT) as the dependent variable. The two-choice optogenetic stimulation and OFT (unilateral stimulation) data were analyzed using three-way repeated measures ANOVAs with virus (ChR2 vs EYFP) as a between-subjects factor, panel (S+ vs S-) and session (day of training) or Laser (ON vs OFF) and turn direction (contralateral vs ipsilateral rotation) as within-subjects factors, and responses or turns as the dependent variable. *Post hoc* Bonferroni’s multiple comparisons tests were performed when ANOVA main effects or interactions were significant (p < 0.05). Mauchly’s sphericity test was used to assess the assumption of sphericity, and the Greenhouse-Geisser correction was applied where necessary (Mauchly’s test p < 0.05).

## Results

### Optogenetic stimulation of the NAc^D1-MSN^-SNr pathway drives reinforcement

Given the established role of NAc D1-MSNs in reinforcement (Cole et al., 2018; Hikida et al., 2010; Lobo et al., 2010), we first investigated whether optogenetic stimulation of the NAc^D1-MSN^-SNr pathway could similarly induce reinforcement. Optic fibers were bilaterally implanted into the SNr of D1-Cre mice that had been microinjected with a Cre-dependent channelrhodopsin 2 (ChR2) (AAV2-Ef1a-FLEX-hChR2(H134R)-EYFP) or EYFP control virus (AAV2-Ef1a-FLEX-EYFP) to enable pathway-specific activation of axon terminals (Fig 1A&B, S1A). We further confirmed that EYFP expression at SNr terminal sites did not overlap with the dopaminergic neurons in the substantia nigra pars compacta (SNc) and the ventral tegmental area (VTA) (Fig S1B). In a real-time place preference (rt-PP) task, mice were first allowed to freely explore a two-chamber apparatus with differing contextual stimuli in each (triangles vs circles) for 15 mins (Pre-test) (Fig 1C). Then, on the next three consecutive days, mice were placed back in the apparatus for 20 mins, which now produced laser stimulation via the optic fibers when mice entered into one of the two chambers (either triangles vs circles, randomized between mice) (Laser tests) (Fig 1C). Mice expressing ChR2, but not EYFP, were found to spend a significantly increased amount of time in the laser-paired chamber in the Laser test sessions compared with in the Pre-test session (Fig 1D&E; Significant virus by session interaction, F_(3,51)_=17.28, p<0.0001), indicating a reinforcing effect of activation of the NAc^D1-MSN^-SNr pathway.

**Fig 1.**
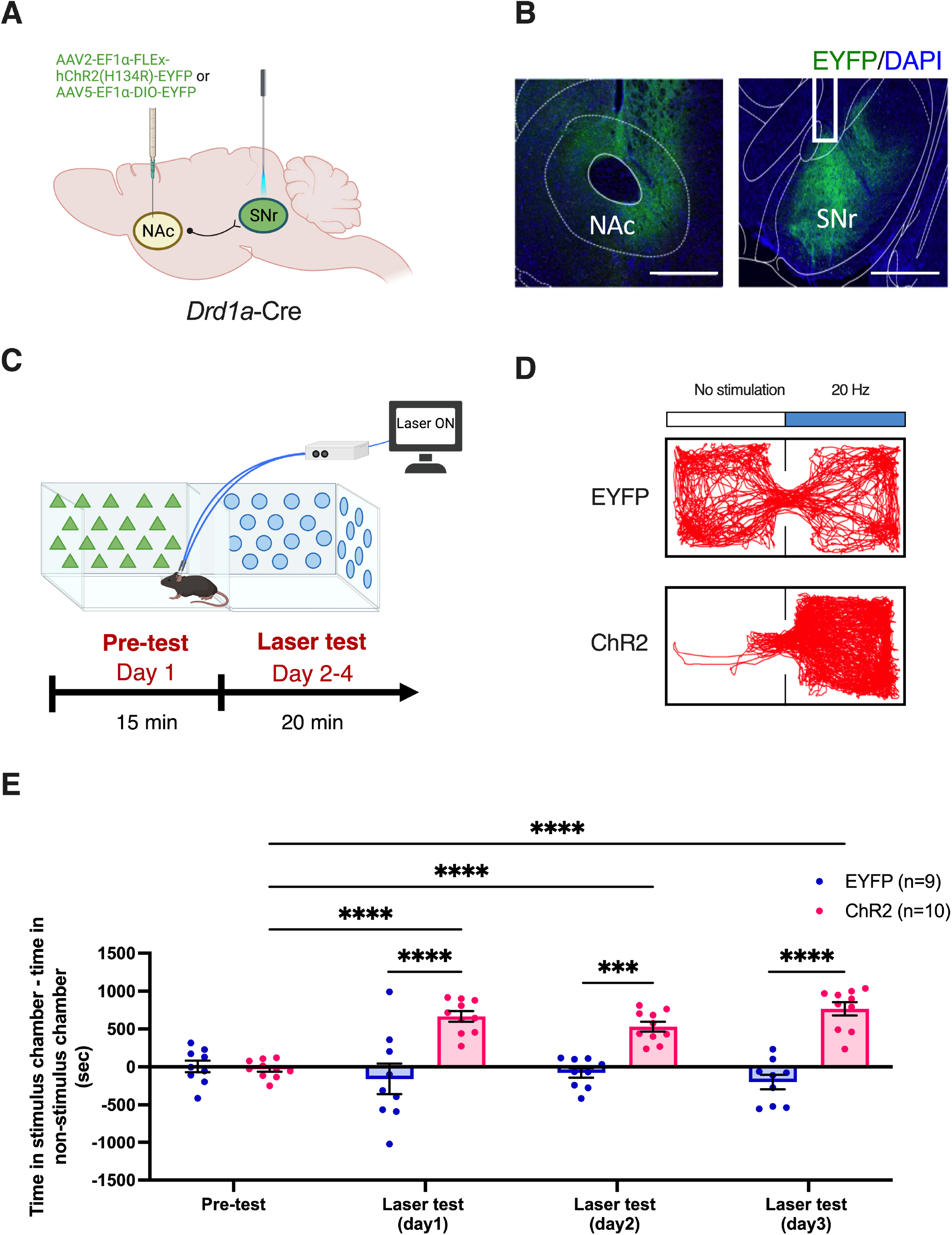
Optogenetic stimulation of the NAc^D1-MSN^-SNr pathway is reinforcing. **(A)** Schematic of viral infusion and optic fiber implantation sites in D1-Cre mice. **(B)** Representative coronal sections of ChR2-EYFP expression (Green) in the nucleus accumbens (NAc) core (Left) and substantia nigra pars reticulata (SNr) (Right) of D1-Cre mice; nuclear marker (DAPI, Blue): scale bar, 500 *µ*m. **(C)** Schematic representation of the real-time place preference (rt-PP) assay and its experimental timeline. **(D)** Representative heatmaps of the time spent in each compartment during the laser-test for two example D1-Cre mice expressing either EYFP or ChR2. **(E)** Time spent in the laser-paired compartment minus time spent in the non-laser-paired chamber in D1-Cre mice expressing EYFP (n=9) or ChR2 (n=10) during pre-test and laser test sessions. Data represent the mean ± SEM, *post hoc* Bonferroni comparisons ****p<0.0001.

### Optogenetic activation of the NAc^D1-MSN^-SNr pathway is sufficient to support instrumental self-stimulation and augments instrumental responding for a liquid reinforcer

To further confirm the functional role of the NAc^D1-MSN^-SNr pathway in reinforcement, we next performed a two-choice schedule of operant self-stimulation. In four consecutive daily 60 min sessions in a touch-screen operant chamber, mice were trained under a fixed-ratio 1 (FR1) schedule of reinforcement to produce a touch response at one of two response panels, one that resulted in a 30-sec delivery of the laser via optic fibers (S+), and the other that resulted in no change (S-) (Fig 2A). Over the course of four test sessions, there was observed to be a significant increase in responding at the S+ panel compared to the S-panel in mice expressing ChR2, but not EYFP (Fig 2B; Significant virus by session by panel interaction, F_(3,51)_=3.19, p<0.05), indicating that activation of NAc D1-MSNs projecting to the SNr is sufficiently reinforcing to sustain instrumental responding. No difference in the latencies to respond at either of the response panels was observed between ChR2- and EYFP-expressing mice (Fig S2A).

**Fig 2.**
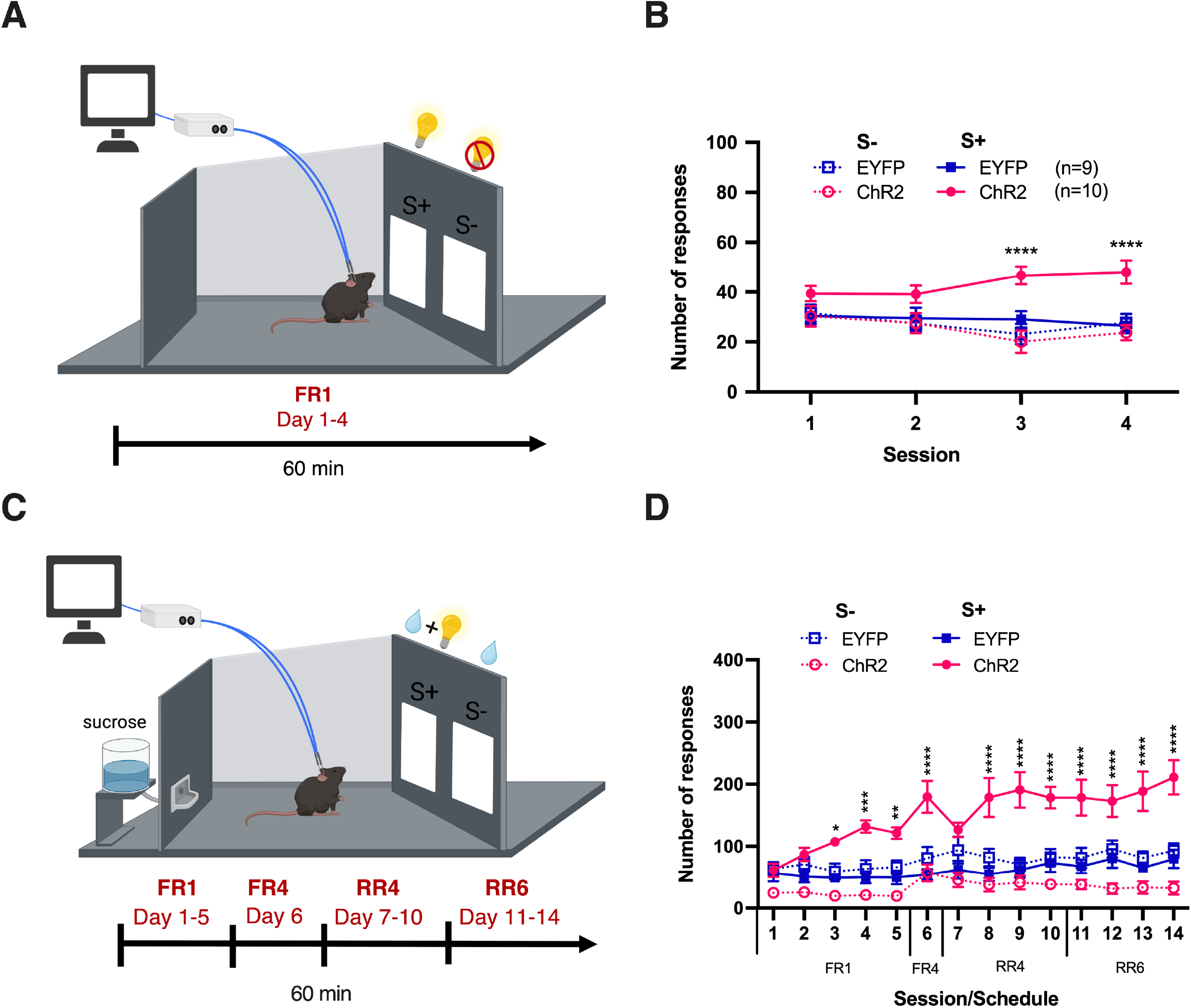
Optogenetic stimulation of NAc^D1-MSN^-SNr pathway supports instrumental self-stimulation and increases instrumental responding for a liquid reinforcer. **(A)** Schematic representation and experimental timeline of the two-choice task in which mice instrumentally respond at two response panels under an FR1 schedule for either self-stimulation by the laser (S+) or no outcome (S-). **(B)** The total number of panel touches in mice receiving optogenetic stimulation of the NAc^D1-MSN^-SNr pathway (ChR2, n=10) and control (EYFP, n=9) mice. **(C)** Schematic representation and experimental timeline of the two-choice task in which mice instrumentally respond at two response panels under an FR1-RR6 schedules for either a liquid reinforcer paired with laser delivery (S+) or the the liquid reinforcer alone (S-). **(D)** The total number of panel touches in the FR1, FR4, RR4, and RR6 sessions of mice receiving optogenetic stimulation of the NAc^D1-MSN^-SNr pathway (ChR2) and control (EYFP) mice. Data represent the mean ± SEM, *post hoc* Bonferroni comparisons *p<0.05, **p<0.01, ***p<0.001, ****p<0.0001.

Next, to investigate whether in addition to be being directly reinforcing, activation of this pathway could modulate the appetitiveness of a liquid reinforcer, we paired delivery of the laser with a response to earn a sucrose reward in a two-choice schedule of reinforcement (Fig 2C). As previously, mice could choose to respond at one of two response panels. However, now a touch response at one randomly assigned panel delivered a sucrose reward alongside a 30 sec laser delivery (S+), while a response at the other panel delivered the sucrose reward by itself (S-). Mice were trained on a previously described schedule of reinforcement with minor modifications (Robinson et al., 2014; Soares-Cunha et al., 2022), that proceeded from FR1 to a random ratio 6 (RR6, a random average of 6 responses needed to produce the outcome) over consecutive days. As with the previous self-stimulation task, it was revealed that ChR2-expressing mice responded a significantly greater amount of times for the S+ panel compared to the S-panel, and that this effect gradually grew stronger across progressing schedules of reinforcement (Fig 2D; Significant virus by session by panel interaction, F_(13,221)_=3.37, P<0.0001). No differences in the latencies to respond at the response panels were observed between ChR2- and EYFP-expressing mice (Fig S2B). However, the latency to collect the liquid reward was observed to be significantly increased when mice expressing ChR2 responded at the S+ panel compared with the S-panel (Fig S2C). It is possible that receipt of the rewarding laser stimulation makes mice less motivated to pick up the liquid reinforcer as they are already receiving a rewarding outcome.

Altogether, the findings of the rt-PP, two-choice self-stimulation, and laser paired with liquid reinforcer two-choice task, indicate that activation of the NAc^D1-MSN^-SNr pathway is reinforcing and able to sustain and augment instrumental responding for the laser itself or a liquid reinforcer, respectively.

### Activation of the NAc^D1-MSN^-SNr pathway increases motor activity in an open field arena

Next, as NAc core D1-MSN activation has previously been associated with augmented motor behavior (Dreher and Jackson, 1989; Plaznik et al., 1989; Zhu et al., 2016), we investigated the effect of stimulation of the NAc^D1-MSN^-SNr pathway on motor activity in an open field arena (Fig 3A). Mice were first habituated to the apparatus for 3 minutes, then underwent 4 alternating 3-min laser off and laser on epochs for a total of 12 mins (Fig 3A). Measurement of motor activity during the laser on vs the laser off epochs revealed that laser stimulation of the axon terminals of NAc core D1-MSNs in the SNr resulted in a significant increase in velocity (Fig 3B; Significant virus by time period interaction F_(11,187)_=3.243, P<0.001, Fig 3C; Significant virus by laser interaction, F_(1,17)_=16.86, P<0.001) and distance moved (Fig 3D&E; Significant virus by laser interaction, F_(1,17)_=14.07, P<0.01).

**Fig 3.**
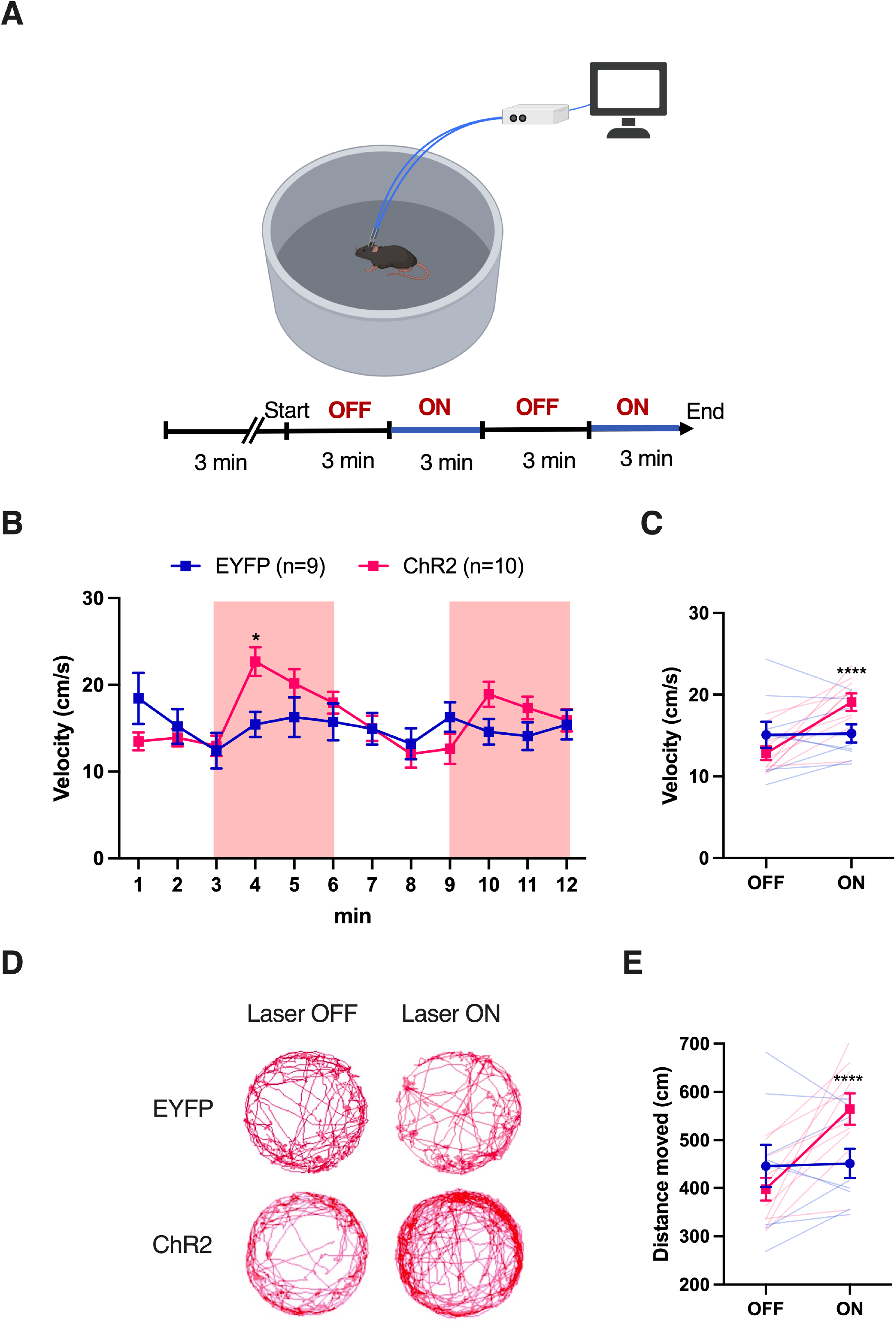
Bilateral stimulation of SNr-projecting NAc D1-MSNs increases locomotor activity. **(A)** Schematic representation and experimental timeline of the open field test in which mice move around a circular chamber during laser OFF and laer ON epochs. **(B)** The velocity of EYFP (n=9) and ChR2 (n=10)-expressing mice during laser OFF and laser ON epochs. Shaded areas indicate the time periods of laser delivery. **(C)** The total distance moved in during laser OFF and laser ON epochs in mice expressing ChR2 or EYFP in the NAc^D1-MSN^-SNr pathway. Data represent the mean ± SEM, *post hoc* Bonferroni comparisons ****p<0.0001.

Finally, we next investigated whether unilateral stimulation is able to bias motor behavior towards the ipsilateral or contralateral direction to the stimulation side. Mice were habituated to the open field apparatus for 3 mins then again underwent alternating 3-min laser off and laser on epochs (Fig 4A). As unilateral stimulation resulting in pronounced turning behavior rather than forward movement, the number of rotations to the ipsilateral or contralateral direction were measured during laser on and off epochs. Stimulation of the NAc^D1-MSN^-SNr pathway resulted in a significant increase in contralateral turns to the stimulation side in mice expressing ChR2, but not EYFP controls (Fig 4B&C, Significant virus by laser by turn direction interaction, F_(1,17)_=6.118, P<0.05 and F_(1,17)_=12.67, P<0.01).

**Fig 4.**
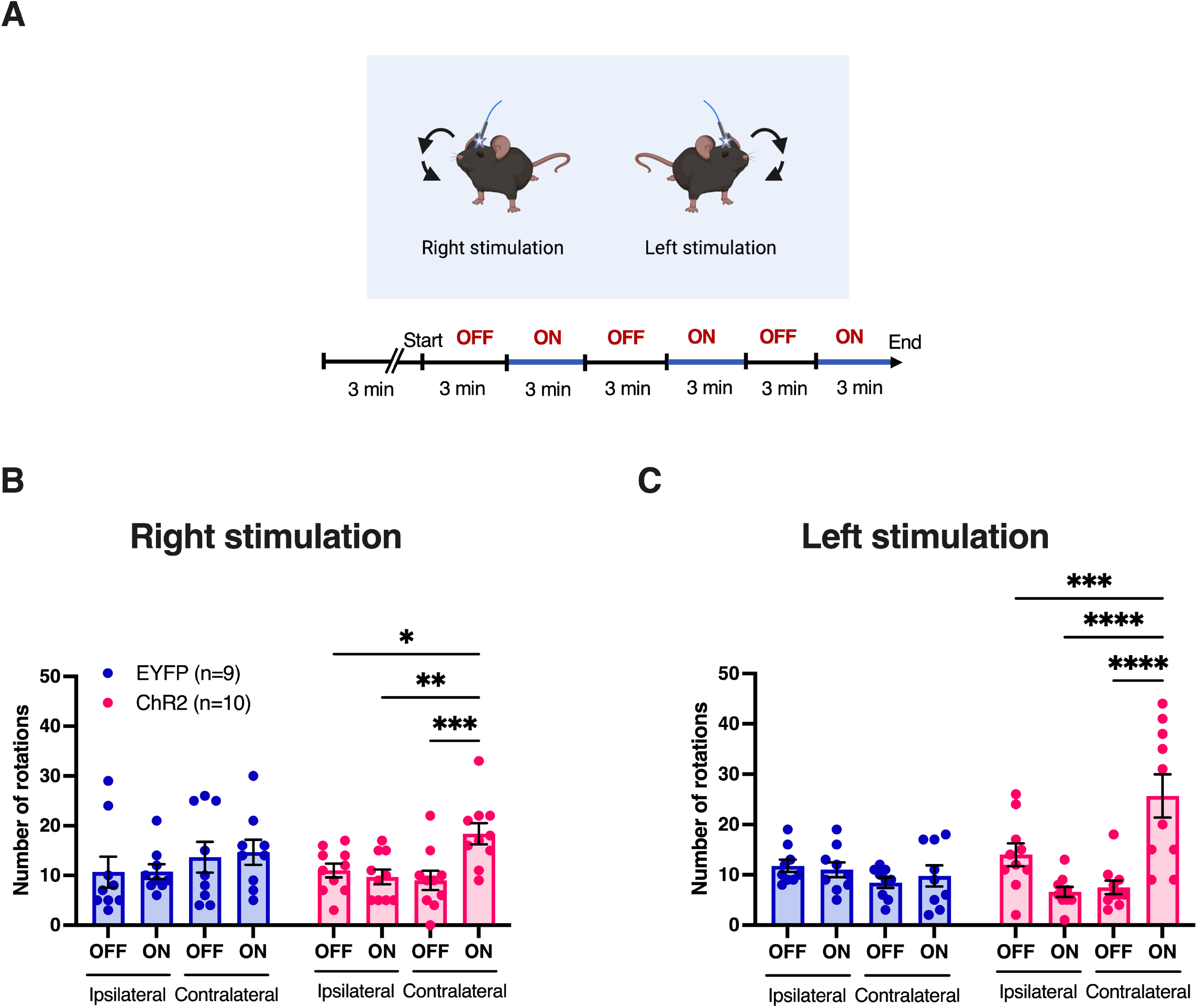
Unilateral stimulation of SNr-projecting NAc D1-MSNs induces contralateral rotations. **(A)** Schematic example of controlateral body rotations in the open field test following unilateral stimulation. **(B)** Total number of ipsilateral and contralateral rotations during right stimulation and **(C)** left stimulation in mice expressing either EYFP (n=9) or ChR2 (n=10). Data represent the mean ± SEM, *post hoc* Bonferroni comparisons *p<0.05, **p<0.01, ***p<0.001, ****p<0.0001.

Together, these findings indicate that activity in NAc core D1-MSNs projecting to the SNr is able to augment motor activity and control turning behavior, suggesting that this pathway may play an important role in motor control.

## Discussion

In this study, we found that activity in NAc core D1-MSNs projecting to the SNr is able to modulate both limbic and motor behaviors. Optogenetic stimulation of the axon terminals of NAc core D1-MSNs in the SNr was directly reinforcing and resulted in an increase in motor activity.

It was observed that mice showed a preference for a location paired with optogenetic stimulation of the NAc^D1-MSN^-SNr pathway and would instrumentally respond in an operant chamber to receive the stimulation alone or the stimulation paired with a liquid reward. These findings support those of previous studies reporting that activation of NAc D1-MSNs is rewarding and able to augment the reinforcing effects of natural or drug rewards (Cole et al., 2018; Hikida et al., 2010; Lobo et al., 2010). Our findings also indicate that the reinforcing effects of cell body stimulation reported in these studies likely occurs via the projection from NAc D1-MSNs to the SNr. Indeed, while NAc core D1-MSNs also project to the VP, a recent study has indicated that optogenetic stimulation of this pathway in mice is aversive rather than reinforcing (Liu et al., 2022). Together these studies suggest a dissociation in the functional roles of NAc core D1-MSNs projecting to the SNr and VP in limbic control.

Our finding that activation of the NAc^D1-MSN^-SNr pathway results in increased motor activity in an open field arena also support the findings of a recent study demonstrating that optogenetic stimulation of the axon terminals of NAc core D1-MSNs in the SNr resulted in increased activity in the motor cortex (M1) (Aoki et al., 2019). It is possible that in our study, increases in velocity following bilateral stimulation and contralateral turning following unilateral stimulation are the result of a facilitation of M1 activity. Indeed, interestingly, it has recently been revealed that the limbic and motor basal ganglia loops may converge within the thalamus via overlapping patterns of innervation from medial and lateral SNr regions which themselves receive input from limbic-related nucleus accumbens and motor-related dorsolateral striatum regions, respectively (Aoki et al., 2019; Foster et al., 2021; Hunnicutt et al., 2014; Macpherson et al., 2021). Importantly, while motor activity was increased, our finding that instrumental responding in the self-stimulation experiments was largely selective to the stimulation-paired panel, rather than a general increase in responses at both panels, suggests that the findings of self-stimulation experiments are not simply the result augmented motor activity.

Interestingly, several psychiatric conditions associated with altered signaling in NAc D1-MSNs are characterized by abnormal limbic- and motor-related behaviors. Indeed, locomotor sensitization, an augmentation of motor activity, is a common early behavioral adaptation to several addictive drugs, including methamphetamine, cocaine, ketamine, alcohol, nicotine, and opioids (Grahame et al., 2000; Ranaldi et al., 2009; Robinson and Berridge, 1993; Strong et al., 2017; Vezina, 2004), and has been reported to coincide with excessive dopamine release in the NAc core and increased activity in NAc D1-MSNs (di Chiara, 2002; van Zessen et al., 2021). Oppositely, motor retardation is a common symptom of both major depression and the depressed phase of bipolar disorder in humans (Buyukdura et al., 2011; Caligiuri and Ellwanger, 2000) and is also reported in chronic social defeat mouse models of depression (Huang et al., 2013; Ota et al., 2018). Accordingly, chronic social defeat in mice is associated with reduced NAc dopamine release and reduced excitatory synaptic input onto NAc D1-MSNs (Francis et al., 2015), an effect that could be predicted to reduce signaling through the NAc^D1-MSN^-SNr pathway. While evidence of a common etiology in the limbic and motor symptoms of psychiatric disorders including depression and substance abuse is yet to be established, our data suggest the NAc^D1-MSN^-SNr pathway may present an attractive target for future investigation.

It should be noted that a limitation of the current study is that only the behavioral effect of NAc^D1-MSN^-SNr pathway stimulation, and not inhibition, was investigated. Given that optogenetic stimulation and inhibition of NAc D1-MSN cell bodies has previously been reported to bidirectionally control reward and aversion, respectively, it could be predicted that optogenetic inhibition of the NAc^D1-MSN^-SNr pathway may similarly result in aversion in rt-PP and self-stimulation studies. This is supported by evidence that knockout of dopamine D1 receptors in mice is sufficient to eliminate prereward anticipatory firing of NAc neurons in a place learning task, and impair intracranial self-stimulation responding (Tran et al., 2005).Interestingly, while chemicogenetic activation of NAc D1-MSNs has been reported to increase locomotor activity, no change in locomotion was observed when NAc D1-MSNs were inhibited (Zhu et al., 2016). In the future, investigation of the behavioral consequences of inhibition of NAc core D1-MSNs projecting the SNr will help to confirm whether this pathway is able to bidirectionally control limbic and motor functions.

Overall, our findings revealed that activity in NAc core D1-MSNs projecting to the SNr was able to influence both reinforcement and motor behavior, indicating a multifunctional role of this pathway. These studies also provide further evidence in support of convergence of limbic and motor basal ganglia circuits, and may help to provide a plausible neural circuit contributing to the frequent cooccurrence of limbic and motor symptoms in psychiatric conditions including depression and substance abuse.

## Author Contributions

SA performed research, analyzed data, and co-wrote the manuscript. TO provided guidance for experiments and co-wrote the manuscript. TM designed the experiments, supervised the project, and co-wrote the manuscript. TH supervised the project and co-wrote the manuscript. All authors read and approved the final manuscript.

## Acknowledgments

We thank Ms. Noriko Otani for her technical assistance. Member of the lab are thanked for constructive feedback throughout this study. Experimental figures created using BioRender.com

## Funding Sources

This work was supported by grants from the Japan Society for the Promotion of Science (JSPS) KAKENHI (JP21K15210 to T.M; JP21H00311, JP21K18557 and JP22H01105, to T.O.; JP18H02542, JP21H00311 and JP22H029440 to T.H.), Japan Agency for Medical Research and Development (AMED) (JP22gm6510012 to T.O.; JP21wm0425010 and 21gm1510006 to T.H.), Salt Science Research Foundation (2146 and 2240 to T.O.; 2137 and 2229 to T.H.), and SENSHIN Medical Research Foundation (to T.H.).

---

**Supplementary Fig S1. Histology of fiber placements. (A)** Representative coronal sections of optic fiber placements in the SNr in EYFP and ChR2-expressing mice. **(B)** Representative image of the EYFP expression in the SNr (Left), TH-positive dopaminergic neurons of SNc and VTA (Middle), and an overlay of both (Right). TH (Red), EYFP (Green), DAPI (Blue). Scale bar, 500 *µ*m.

**Supplementary Fig S2. Optogenetic stimulation of the NAc**^**D1-MSN**^**-SNr pathway during two-choice optogenetic stimulation tasks does not affect the latency to make responses, but is able to increase the latency to collect a liquid reward. (A)** The latency to make a response at the S+ or S-panel in the two-choice optogenetic self-stimulation task in mice expressing either EYFP (n=9) or ChR2 (n=10). **(B)** The latency to make a response at the S+ or S-panel in the two-choice laser & liquid reinforcer task. **(C)** The latency to collect the liquid reward in the two-choice laser & liquid reinforcer task. ChR2 mice showed an increased latency to collect the liquid reward following an S+ response. Data represent the mean ± SEM, *post hoc* Bonferroni comparisons **p<0.01.

**Supplementary Table S1**. Detailed statistical analyses of all experiments included in the manuscript.

## Notes

### Competing Interest Statement

The authors have declared no competing interest.

