## Supplementary figures and images for "Dual roles for nucleus accumbens core dopamine D1-expressing neurons projecting to the substantia nigra pars reticulata in limbic and motor control"

### Supplementary Figure 1

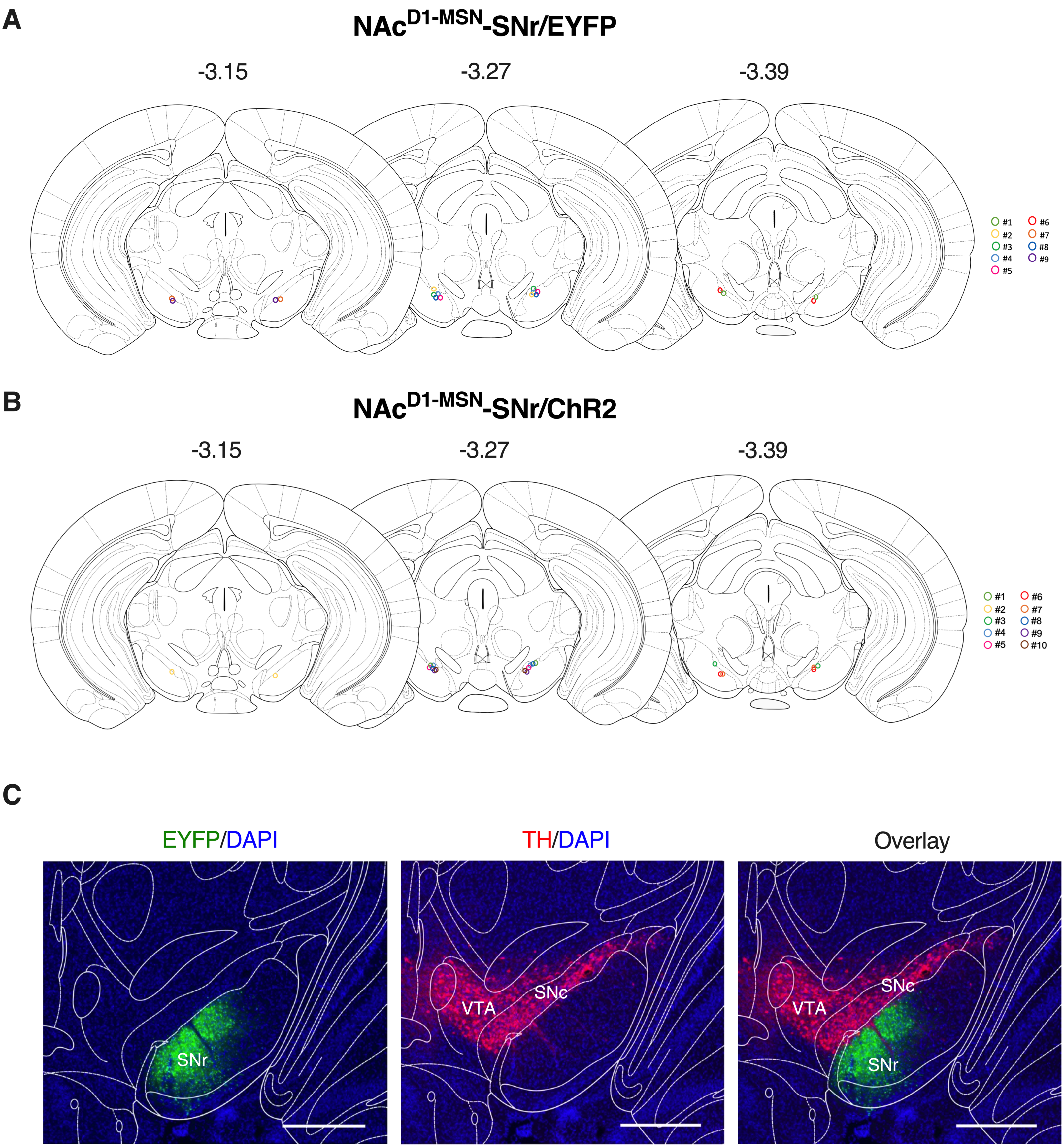

### Supplementary Figure 2

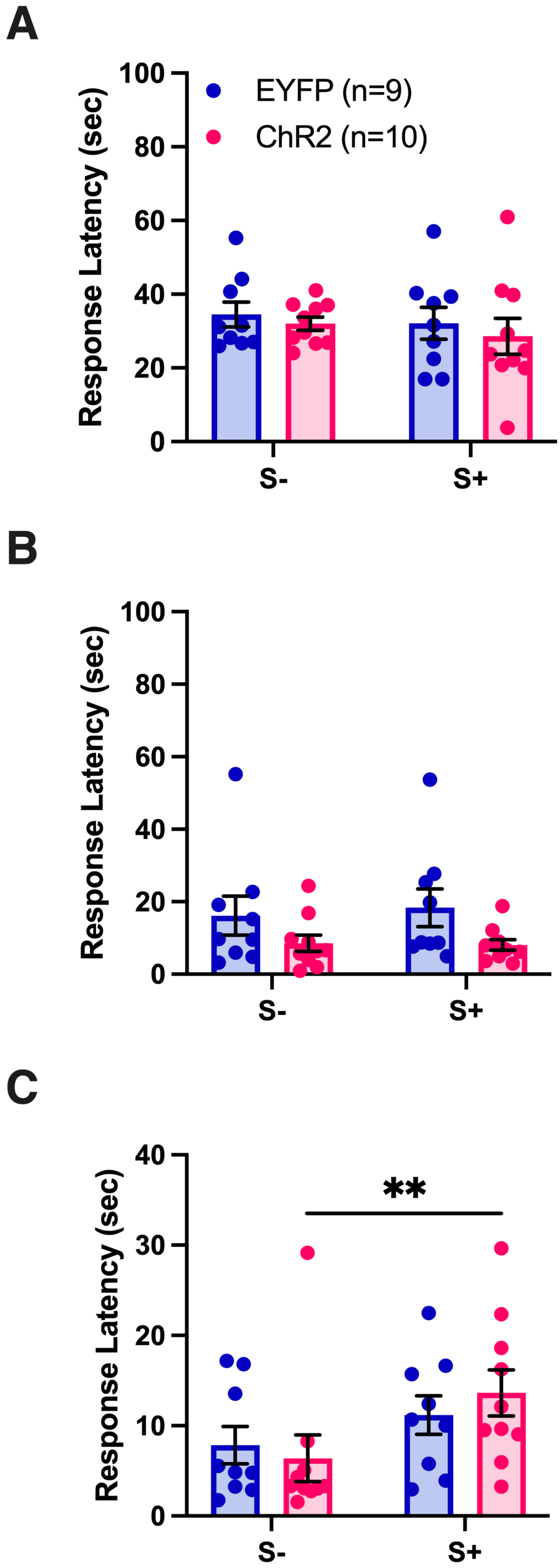
