## Supplementary Table 1 for "Dual roles for nucleus accumbens core dopamine D1-expressing neurons projecting to the substantia nigra pars reticulata in limbic and motor control"

| Figure | | Description | N-Number | Statistic | P-value | Post-hoc | Post-hoc P-value |
|---|---|---|---|---|---|---|---|
| 1 | E | rt-PP, the preference time spend in the laser-paired chamber (sec) | D1-SNr/ChR2 10 mice<br>D1-SNr/EYFP 9 mice | 2-way RM Anova<br><br>virus F (1,17) = 35.52<br>session F (3,51) = 6.361<br>virus x session F (3, 51) = 17.28 | p<0.0001<br>p<0.001<br>p<0.0001 | Bonferroni | ChR2, Pre-test vs Laser test (day1), ****p<0.0001<br>ChR2, Pre-test vs Laser test (day2), ****p<0.0001<br>ChR2, Pre-test vs Laser test (day3), ****p<0.0001<br>ChR2 vs EYFP, Laser test (day1), ****p<0.0001<br>ChR2 vs EYFP, Laser test (day2), ***p<0.001<br>ChR2 vs EYFP, Laser test (day3), ****p<0.0001 |
| 2 | B | Two choice optogenetic stimulation, number of responses (day1-4) | D1-SNr/ChR2 10 mice<br>D1-SNr/EYFP 9 mice | 3-way RM Anova<br><br>virus F (1,17) = 2.511<br>panel F (1,17) = 20.87<br>session F (3,51) = 1.301<br>virus x panel F (1,17) = 15.20<br>virus x session F (3,51) = 0.9290<br>panel x session F (3,51) = 5.905<br>virus x panel x session F (3,51) = 3.193 | p=0.1315<br>p<0.001<br>p=0.2844<br>p<0.01<br>p=0.4335<br>p<0.01<br>p<0.05 | Bonferroni | ChR2, S- vs S+, ****p<0.0001 (day3)<br>ChR2 (S+) vs EYFP (S-), ns (day3)<br>ChR2 vs EYFP, S+, p<0.01 (day3)<br>ChR2, S- vs S+, ****p<0.0001 (day4)<br>ChR2 (S+) vs EYFP (S-), *p<0.05 (day4)<br>ChR2 vs EYFP, S+, p<0.01 (day4) |
| | D | Two choice optogenetic stimulation paired with a liquid reinforcer, number of responses (day1-14) | | 3-way RM Anova<br><br>virus F (1,17) = 5.829<br>panel F (1,17) = 22.57<br>session F (13,221) = 7.238<br>virus x panel F (1,17) = 39.59<br>virus x session F (13,221) = 2.574<br>panel x session F (13,221) = 3.504<br>virus x panel x session F (13,221) = 3.365 | p<0.05<br>p<0.001<br>p<0.0001<br>p<0.0001<br>p<0.01<br>p<0.0001<br>p<0.0001 | Bonferroni | ChR2, S- vs S+, *p<0.05 (day3)<br>ChR2, S- vs S+, ***p<0.001 (day4)<br>ChR2, S- vs S+, **p<0.01 (day5)<br>ChR2, S- vs S+, ****p<0.0001 (day6)<br>ChR2, S- vs S+, ****p<0.0001 (day8)<br>ChR2, S- vs S+, ****p<0.0001 (day9)<br>ChR2, S- vs S+, ****p<0.0001 (day10)<br>ChR2, S- vs S+, ****p<0.0001 (day11)<br>ChR2, S- vs S+, ****p<0.0001 (day12)<br>ChR2, S- vs S+, ****p<0.0001 (day13)<br>ChR2, S- vs S+, ****p<0.0001 (day14) |
| 3 | B | OFT, bilateral stimulation, velocity (cm/s) per min | D1-SNr/ChR2 10 mice<br>D1-SNr/EYFP 9 mice | 2-way RM Anova<br><br>virus F (1,17) = 0.3711<br>time period F (11,187) = 4.090<br>virus x laser F (11, 187) = 3.243 | p=0.5505<br>p<0.0001<br>p<0.001 | Bonferroni | ChR2 vs EYFP, min 4, *p<0.05 |
| | C | OFT, bilateral stimulation, velocity (cm/s) | | 2-way RM Anova<br><br>virus F (1,17) = 0.2831<br>laser F (1,17) = 18.90<br>virus x laser F (1, 17) = 16.86 | p=0.6016<br>p<0.001<br>p<0.001 | Bonferroni | ChR2, OFF vs ON , ****p<0.0001<br>EYFP, OFF vs ON , ns |
| | E | OFT, bilateral stimulation, distance moved (cm) | | 2-way RM Anova<br><br>virus F (1,17) = 0.5944<br>laser F (1,17) = 15.97<br>virus x laser F (1, 17) = 14.07 | p=0.4513<br>p<0.001<br>p<0.01 | Bonferroni | ChR2, OFF vs ON , ****p<0.0001<br>EYFP, OFF vs ON , ns<br>ChR2 vs EYFP, ON, *p<0.05 |
| 4 | B | OFT, unilateral stimulation (Right hemisphere), number of rotations | D1-SNr/ChR2 10 mice<br>D1-SNr/EYFP 9 mice | 3-way RM Anova<br><br>virus F (1,17) = 0.03149<br>laser F (1,17) = 5.475<br>turn direction F (1,17) = 5.194<br>virus x laser F (1,17) = 3.152<br>virus x turn direction F (1,17) = 0.001004<br>laser x turn direction F (1,17) = 8.536<br>virus x laser x turn direction F (1,17) = 6.118 | p=0.8612<br>p<0.05<br>p<0.05<br>p=0.0937<br>p=0.9751<br>p<0.01<br>p<0.05 | Bonferroni | ChR2, ipsilateral vs contralateral, ON, *p<0.05<br>ChR2, contralateral rotations, OFF vs ON , **p<0.01 |
| | C | OFT, unilateral stimulation (Left hemisphere), number of rotations | | 3-way RM Anova<br><br>virus F (1,17) = 2.878<br>laser F (1,17) = 8.348<br>turn direction F (1,17) = 1.886<br>virus x laser F (1,17) = 6.794<br>virus x turn direction F (1,17) = 8.579<br>laser x turn direction F (1,17) = 17.63<br>virus x laser x turn direction F (1,17) = 12.67 | p=0.1080<br>p<0.05<br>p<0.0001<br>p<0.05<br>p<0.01<br>p<0.001<br>p<0.01 | Bonferroni | ChR2, ipsilateral vs contralateral, ON, ****p<0.0001<br>ChR2, contralateral rotations, OFF vs ON , ****p<0.0001<br>ChR2 vs EYFP, contralateral rotations, ON, ****p<0.0001 |
| S2 | A | Two choice optogenetic stimulation, response latency (sec) | D1-SNr/ChR2 10 mice<br>D1-SNr/EYFP 9 mice | 2-way RM Anova<br><br>virus F (1,17) = 0.4192<br>laser F (1,17) = 1.249<br>virus x laser F (1, 17) = 0.04124 | p=0.5260<br>p=0.2792<br>p=0.8415 | N/A | N/A |
| | B | Two choice optogenetic stimulation paired with a liquid reinforcer, response latency (sec) | | 2-way RM Anova<br><br>virus F (1,17) = 2.812<br>laser F (1,17) = 1.371<br>virus x laser F (1, 17) = 3.379 | p=0.1118<br>p=0.2577<br>p=0.0836 | N/A | N/A |
| | C | Two choice optogenetic stimulation paired with a liquid reinforcer, reward correction latency (sec) | | 2-way RM Anova<br><br>virus F (1,17) = 0.02591<br>laser F (1,17) = 18.52<br>virus x laser F (1, 17) = 2.519 | p=0.8740<br>p<0.001<br>p=0.1309 | Bonferroni | ChR2, S- vs S+ , **p<0.01<br>EYFP, S- vs S+ , ns<br>ChR2 vs EYFP, S+, ns |
